# Loss of embryonically-derived Kupffer cells during hypercholesterolemia accelerates atherosclerosis development

**DOI:** 10.1101/2023.09.26.559586

**Authors:** Rebecca Fima, Sébastien Dussaud, Cheïma Benbida, Margault Blanchet, François Lanthiez, Lucie Poupel, Claudia Brambilla, Martine Moreau, Alexandre Boissonnas, Emmanuel L. Gautier, Thierry Huby

## Abstract

Hypercholesterolemia is a major risk factor for atherosclerosis and associated cardiovascular diseases. The liver plays a key role in the regulation of plasma cholesterol levels and hosts a large population of tissue-resident macrophages known as Kupffer cells (KCs). KCs are located in the hepatic sinusoids where they ensure key functions including blood immune surveillance. However, how KCs homeostasis is affected by the build-up of cholesterol-rich lipoproteins that occurs in the circulation during hypercholesterolemia remains unknown. Here, we found that embryo-derived KCs (EmKCs) accumulated large amounts of lipoprotein-derived cholesterol, in part through the scavenger receptor CD36, and massively expanded early after the induction of hypercholesterolemia. After this rapid adaptive response, EmKCs exhibited mitochondrial oxidative stress and their numbers gradually diminished while monocyte-derived KCs (MoKCs) with reduced cholesterol-loading capacities seeded the KC pool. Decreased proportion of EmKCs in the KC pool enhanced liver cholesterol content and exacerbated hypercholesterolemia, leading to accelerated atherosclerotic plaque development. Together, our data reveal that KC homeostasis is perturbed during hypercholesterolemia, which in turn alters the control of plasma cholesterol levels and increases atherosclerosis.

## Main

Atherosclerotic cardiovascular disease (ASCVD) and its clinical outcomes such as myocardial infarction are a leading cause of morbidity and mortality worldwide. Multiple lines of evidence have established that hypercholesterolemia, characterized by high concentrations of cholesterol-rich low-density lipoproteins (LDL)^1^ and their oxidatively damaged forms (oxLDL)^2^, is a leading risk factor in ASCVD. While the liver and intestine are the major organs regulating plasma cholesterol levels, several lines of evidence suggest the immune system could also participate. Thus, reducing^3,4^ or increasing^5^ tissue-resident macrophage (trMac) numbers associated with plasma cholesterol elevation or reduction, respectively. Yet, whether a specific trMac population is primarily involved remains undefined. Though, these models had in common a modulation in the hepatic trMacs, Kupffer cells (KCs), arguing for their potential implication in the control of cholesterolemia.

Located in the hepatic sinusoids, KCs are the largest trMac population in direct contact with the blood and thus with circulating lipoproteins. They originate from various progenitor waves during embryogenesis and maintain by self-renewal independently from circulating monocytes in the steady-state adult liver^6–8^. However, when the embryo-derived KC (EmKC) network is altered during non-alcoholic steatohepatitis (NASH)^9–12^, excessive intravascular hemolysis^13^, bacterial^14^ or parasitic^15^ infections, blood monocytes can be recruited to the liver and differentiate into KCs. Analysis of how these monocyte-derived KCs (MoKCs) engraft the liver tissue has revealed that KCs closely interact with liver sinusoidal endothelial cells (LSECs), hepatic stellate cells (HSCs) and hepatocytes in order to acquire their tissue-imprinted signature as well as the signals allowing them to self-maintain^16,17^. Aside their undisputable role as an intravascular immune barrier that constantly filters the blood for pathogens^18,19^, but also for damaged red blood cells^13^, KCs have been recently endowed with activities influencing the liver response to metabolic diseases such as obesity^20^ or NASH^10^.

Here, we evaluated whether KCs homeostasis was altered in the context of hypercholesterolemia using cholesterol-fed mice deficient for the LDL receptor gene (*Ldlr*^-/-^).

Our findings reveal a two-phase adaptation of the KC pool to plasma cholesterol elevation. In a first stage, EmKCs adapt to this environmental change by loading large amount of cholesterol and increasing their tissue density. Then, in a second phase, EmKCs are gradually lost over weeks and replaced by MoKCs exhibiting a diminished ability to uptake lipoprotein-derived cholesterol. To infer the functional consequences of EmKCs replacement by MoKCs, we specifically deleted EmKCs. Replacement of EmKCs by MoKCs in hypercholesterolemic mice resulted in perturbed hepatic cholesterol homeostasis with decreased activation of the retinoid X receptors (RXR) / liver X receptors (LXR) pathway that associated with increased cholesterol content in the liver tissue and in plasma. Overall, our results support a role for EmKCs in regulating hepatic metabolic adaptation to hypercholesterolemia.

## Results

### Embryo-derived Kupffer cells markedly increased shortly after the induction of hypercholesterolemia

To characterize the potential changes in hepatic leukocyte populations that occur rapidly after the induction of hypercholesterolemia, *Ldlr*^-/-^ mice were subjected to a chow diet enriched with 1% cholesterol (HC diet). After 4 days of HC diet, cholesterolemia raised by 4 times, reaching approximately 1000 mg/dL (Fig. 1A). We then performed a flow cytometry analysis of common hepatic leukocyte populations in livers perfused *in situ* with collagenase. Of note, *in situ* perfusion was critical to retrieve KCs under hypercholesterolemia as classical digestion protocols failed. A t-distributed stochastic neighbor embedding (tSNE) analysis of the cytometry data revealed no major changes between chow and HC diet conditions, except for KCs. KCs, identified by their classical marker VSIG4, distributed very differently between conditions in the tSNE projection (Fig. 1B). To further analyze the impact of short-term HC diet feeding on KCs, RNA sequencing was performed on KCs isolated from mice fed a chow or a HC diet for 4 days. This revealed marked changes in gene expression upon HC diet (Fig. 1C). KEGG pathway analysis on genes up-regulated in the HC-fed condition showed terms associated with lysosomes and cholesterol metabolism (Fig. 1D). Highlighted pathways were also suggestive of intense metabolic activities (metabolism of xenobiotics, arginine biosynthesis, drug metabolism, ribosome). Interrogation of the GSEA Molecular Signatures Database (MSigDB) confirmed such activities (Xenobiotic, Mtorc1 signaling, Glycolysis) and also revealed the induction of genes involved in cell cycling (G2m checkpoint, Mitotic spindle) (Fig. 1D). We then assessed whether the enrichment in pathways linked to cell proliferation was associated with increased KC numbers. Our previous studies have demonstrated that all CLEC2^hi^ leukocytes observed by flow cytometry in the healthy or diseased liver identified KCs, including MoKCs that lack TIMD4 expression^10^. After 4 days of hypercholesterolemia, CD45^+^ CLEC2^+^ KCs displayed, similar to the chow condition, homogeneous cell surface expression for the EmKCs markers CLEC4F, VSIG4 and TIMD4^10,21^ (Fig. 1E). Thus, all KCs remain of embryonic origin after 4 days of HC diet and EmKCs absolute numbers increased by 2-3-fold in the HC-fed condition (Fig. 1F). Increased KC density was confirmed by immunofluorescence microscopy of frozen sections using E-CADHERIN staining to demarcate periportal regions^22^ and CLEC4F staining to identify KCs (Fig. 1G). This led us to assess the proportion of KCs expressing the proliferation marker KI-67. The results showed approximately a 4-fold increase in KI-67^+^ EmKCs in mice fed the HC diet for 4 days as compared to mice fed the chow diet (Fig. 1H). A kinetic analysis showed that increased EmKCs proliferation was already noticeable as soon as 3 days after the start of HC diet feeding (Extended data Fig. 1). Thus, as suggested by our RNA-Seq data, HC diet-induced hypercholesterolemia increased EmKCs proliferation and density in the liver of HC diet-fed animals. We next wondered what drove EmKCs proliferation and focused on colony-stimulating factor 1 (CSF1) that is known to play a critical role in tissue macrophage maintenance and proliferation^16,23^. RT-qPCR measurement of *Csf1* mRNA expression showed its progressive elevation from day 2 to day 4 in the livers of *Ldlr*^-/-^ mice fed the HC diet (Fig. 1I). By contrast, *Il34* mRNA, which encode IL-34 the other known ligand of CSF1 receptor (CSF1R), was only minimally increased (data not shown). Administration of the CSF1R inhibitor PLX3397 to *Ldlr*^-/-^ mice during the first 2-days of HC diet feeding fully blocked EmKCs proliferation and increase in numbers (Fig. 1J). Thus, EmKCs proliferate and expand shortly after the induction of hypercholesterolemia in a CSF1R signaling-dependent manner.

**Fig. 1.**
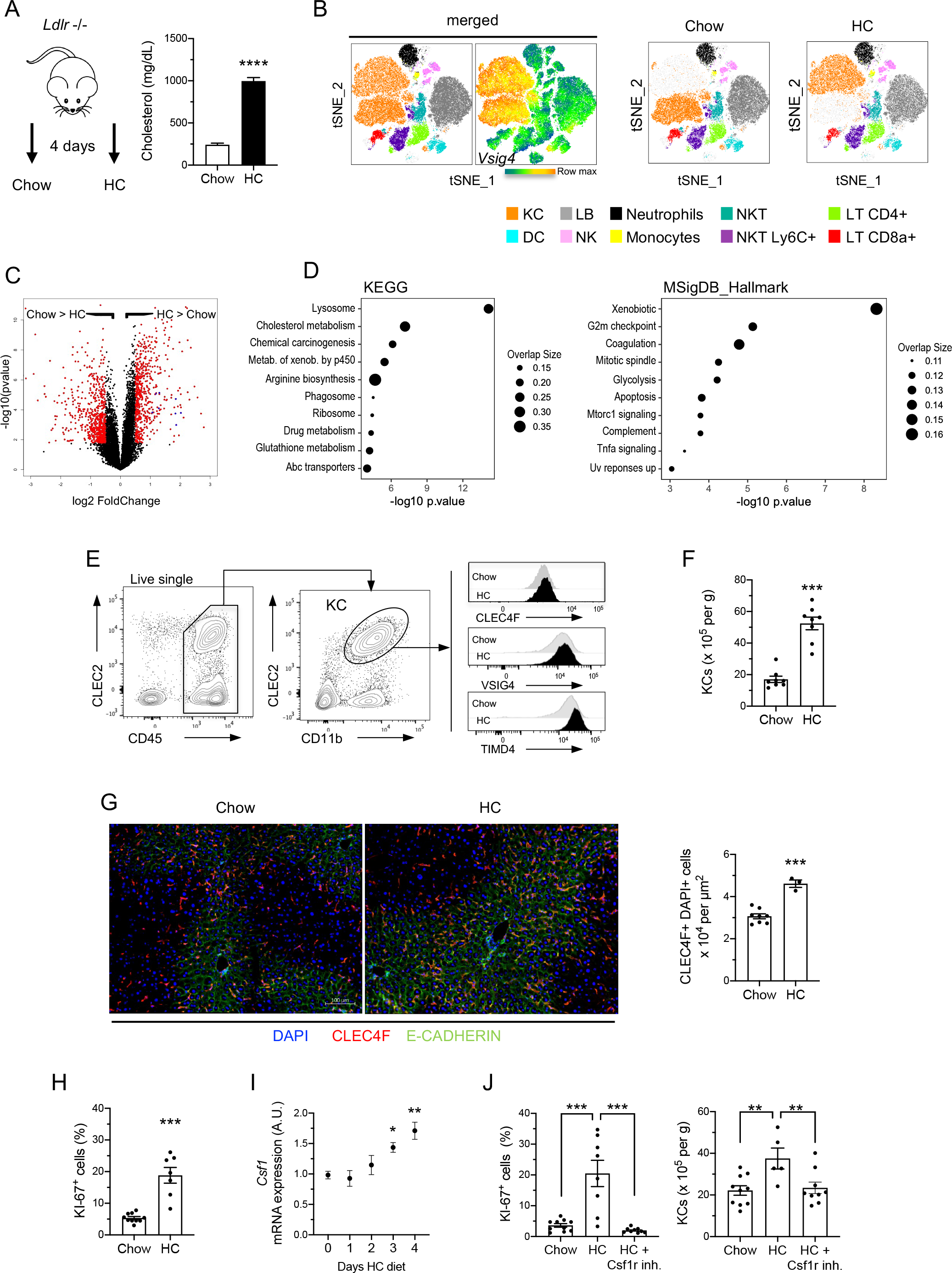
Kupffer cell pool rapidly expands in response to induction of hypercholesterolemia. (A) Plasma total cholesterol concentration in *Ldlr*−/− male mice fed a chow diet (n=8 mice) or chow diet supplemented with 1% cholesterol (HC) (n=7 mice) for 4 days. (B) tSNE projections of identified leukocytes by flow cytometry in livers of chow-fed or HC-fed *Ldlr*−/− mice for 4 days. High expression of VSIG4 identifies KC population. (C) Volcano plot depicting differentially expressed genes (fold change ≥1.3 and adjusted p-value ≥ 0.05 are shown in red) in sorted KCs (CD45^+^CLEC2^+^TIMD4^+^CD31^−^) from livers of chow-fed or HC-fed *Ldl*r−/− mice for 4 days. (D) Pathways enriched in KCs in response to hypercholesterolemia induction. (E) Flow cytometry analysis of KCs and histograms showing the level of expression of the common KC markers CLEC4F, VSIG4 and TIMD4 in both chow- and HC-fed conditions. (F) KC density in livers of chow-(n=8 mice) and HC-fed (n=8 mice) *Ldlr*−/− male mice for 4 days, as determined by flow cytometry. (G) Representative images of epifluorescence microscopy of livers of chow- and HC-fed (4 days) *Ldl*r−/− male mice showing expression of CLEC4F (red). E-CADHERIN (green) staining highlights the preferential localization of KCs in periportal regions. Nuclei (DAPI) are shown in blue. Quantification of CLEC4F+ and DAPI+ cells density confirmed increased values in HC-fed (4 days, n=3) as compared to chow-fed (n=8) *Ldlr*−/− mice. (H) Percentage of KI-67^+^ cells among CLEC2^+^ KCs of chow (n=10) or HC-fed (n=8) *Ldlr*−/− male mice, confirming increased KCs proliferation following induction of hypercholesterolemia. (I) Fold change in *Csf1* gene expression in livers of *Ldlr*−/− male mice fed HC diet for the indicated days (n=12, 3, 7, 6 and 10 mice for day 0, 1, 2, 3 and 4, respectively). Data are expressed relative to day 0 and p values correspond to statistical differences versus day 0. (J) Percentage of KI-67^+^ KCs and KC numbers in *Ldlr*−/− female mice fed a chow or HC diet for 3 days and treated with PLX3397 (CSF1R inhibitor) or vehicle.

### Massive cellular accumulation of cholesterol occurs in Kupffer cells following induction of hypercholesterolemia

We next aimed to further characterize the cellular changes occurring in EmKCs shortly after the induction of hypercholesterolemia (4 days). As reported above, genes associated with cholesterol metabolism were enriched in KCs isolated from hypercholesteremic animals (Fig 1D). Thus, we wondered if genes involved in cholesterol biosynthesis were regulated in KCs after the induction of hypercholesteremia. We performed Gene Set Enrichment Analysis (GSEA) using a gene dataset that include all the major genes involved in cholesterol synthesis. The analysis revealed a clear downregulation of this pathway in the HC-fed condition (Fig 2A). Such expression profile is usually observed in cholesterol-loaded cells, including foamy macrophages^24^. Foamy macrophages are typically found in atherosclerotic lesions and transcriptomic profiles of intimal foamy and non-foamy macrophages were recently reported^25^. We thus conducted GSEA using the 250 most up-regulated genes in foamy versus non-foamy macrophages and revealed a specific enrichment of these genes in EmKCs isolated from HC-fed *Ldlr*^-/-^ mice (Fig. 2A). Foamy macrophages found in atherosclerotic plaques were previously characterized by their high granularity and elevated intracellular lipid levels as revealed by bodipy 493/503 staining^25^. Flow cytometry analyses also showed that hypercholesterolemia specifically elevated granularity (SSC-A parameter) in KCs (Fig. 2B) that associated with higher bodipy staining (Fig. 2C). Thus, KCs accumulate intracellular lipid shortly after the induction of hypercholesterolemia. Targeted lipidomic showed an increase in both free cholesterol (4-fold) and cholesteryl-esters (CE) (15-fold) in KCs isolated from mice fed the HC diet as compared to chow-fed controls (Fig. 2D). All CE species quantified were found elevated in cholesterol-loaded KCs with CE derived from the monounsaturated fatty acid oleate (CE 18:1) being the most prevailing form (Fig. 2D). We then used two-photon laser scanning microscopy combined with Coherent anti-Stokes Raman Spectroscopy (CARS) to detect lipids *in situ* in the liver of *Ldlr*^-/-^ mice fed chow and HC diet for 4 days. We observed the presence of numerous lipid droplets in KCs of hypercholesterolemic mice (Fig. 2E) and, as expected, larger lipid droplets were also detectable in hepatocytes under these conditions (Fig. 2E). Lipidomic analysis of the liver revealed that hepatic cholesterol content raised only by 2-fold after 4 days of hypercholesterolemia, and primarily in the form of CE (Extended data Fig. 2A). Considering both KC numbers and cholesterol-loading increased following the induction of hypercholesterolemia, we estimated a 25-fold increase in the amount of cholesterol stored by the KC population upon HC feeding (Extended data Fig. 2B). Nevertheless, despite this important increase, the amount of cholesterol stored in KCs was only a marginal fraction of the total liver cholesterol content as it was estimated to represent only 0.25%.

**Fig. 2.**
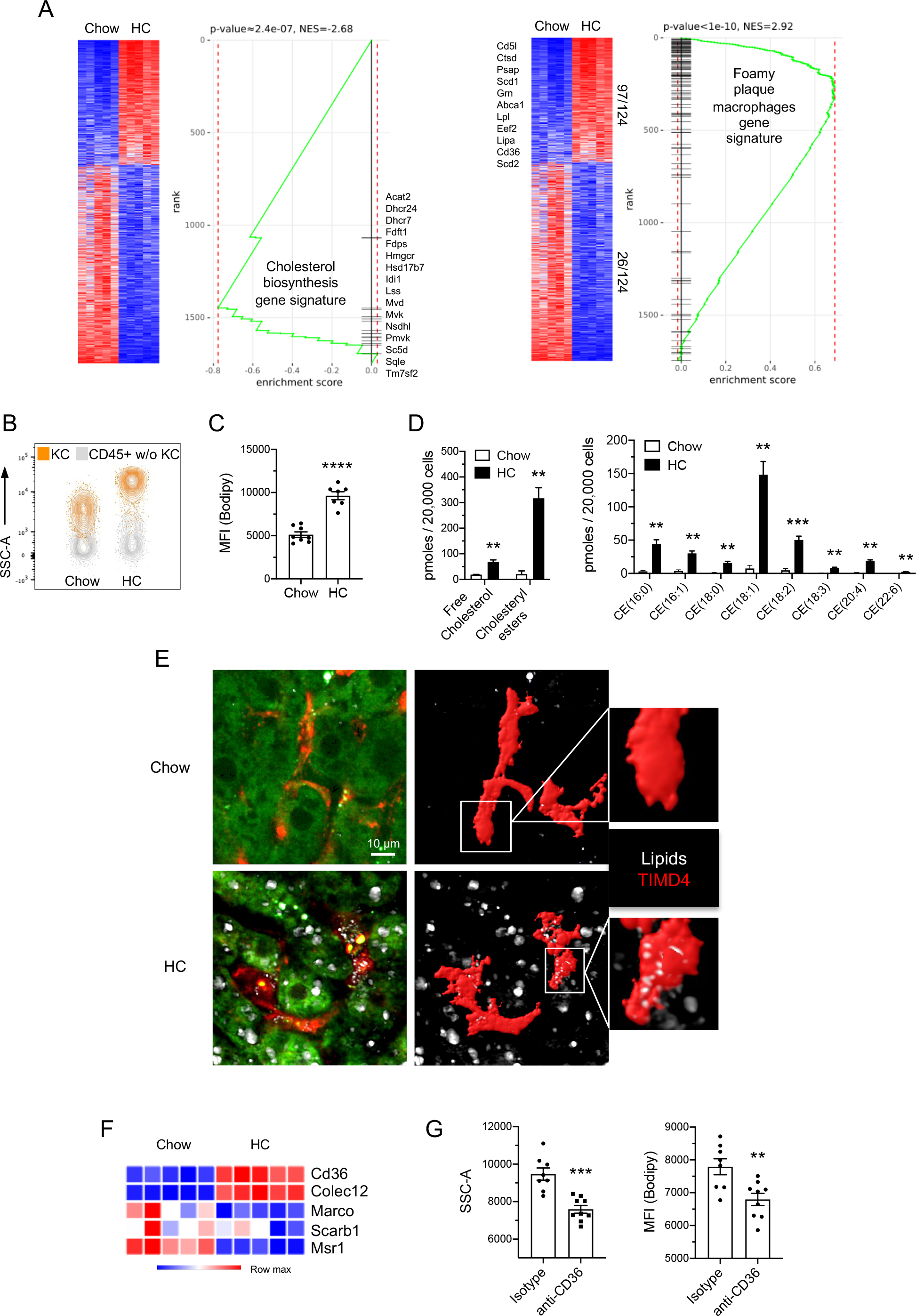
Hypercholesterolemia generates cholesterol-loaded foamy KCs. (A) Gene Set Enrichment analysis (GSEA) of the reactome_cholesterol_biosynthesis and foamy plaque macrophages gene signatures in KC from chow- and HC-fed (4 days) *Ldlr* −/− mice using the Phantasus software. The foamy plaque macrophages gene signature was generated from a previously published dataset (GSE116239). (B) KCs in HC-fed mice exhibit high granularity (SSC-A^hi^) and (C) strongly stain for lipids (bodipy). (D) Changes in cholesterol content, and in cholesteryl-esters species in KCs sorted from chow- and HC-fed (4 days) *Ldlr* −/− mice. (E) Two-photon laser scanning microscopy images of the liver of chow and HC-fed (4 days) *Ldlr* −/− mice. Numerous lipid droplets (white spots) are detected by Coherent anti-Stokes Raman Spectroscopy (CARS) in TIMD4^+^ KCs (red; 3D reconstruction by mask rendering on the right images) and in the hepatocytes (autofluorescence in green) of HC-fed mice as compared to the chow condition. (F) Heatmap generated from the RNA-Seq depicting the level of expression of scavenger receptors in KCs from chow- and HC-fed (4 days) *Ldlr* −/− mice. (G) Changes in the granularity (SSC-A) and lipid content (bodipy) of KCs in mice fed HC diet overnight and injected *in vivo* either with anti-CD36 blocking antibodies or isotype control antibodies.

Considering the phagocytic nature of KCs, we hypothesized that cholesterol loading in KCs would result from the uptake of modified forms of LDL generated upon hypercholesterolemia, as it occurs in atherosclerotic foamy macrophages. Among the different scavenger receptors involved in modified LDLs uptake, *Cd36*^26^ and *Colec12*^27^ were found to be up-regulated in KCs from HC-fed animals (Fig. 2F). To test the potential involvement of CD36, we administered a CD36 blocking mAbs^28^ or its isotype control to *Ldlr*^-/-^ fed the HC diet for 1 day. Blocking CD36 led to decreased KC granularity as well as bodipy staining (Fig. 2G). Thus, cholesterol loading is, at least partially, dependent on CD36.

We next sought to exclude the possibility that the changes observed in KCs were derived from other mechanisms than the hypercholesterolemic state, such as dietary cholesterol-induced gut microbiota dysbiosis^29^. To do so, *Ldlr*^-/-^ mice were treated with the selective cholesterol absorption inhibitor ezetimibe while given cholesterol in the diet. A pre-treatment period with ezetimibe before addition of cholesterol was set up to ensure full activity of the inhibitor (Fig. 3A). Ezetimibe treatment fully blocked plasma cholesterol elevation when mice were switched to HC diet (Fig. 3B). As a result, KCs proliferative response (Fig. 3C), expansion (Fig. 3D) and lipid loading (Fig. 3E) were abolished, demonstrating that induction of hypercholesterolemia was the only triggering factor.

**Fig. 3.**
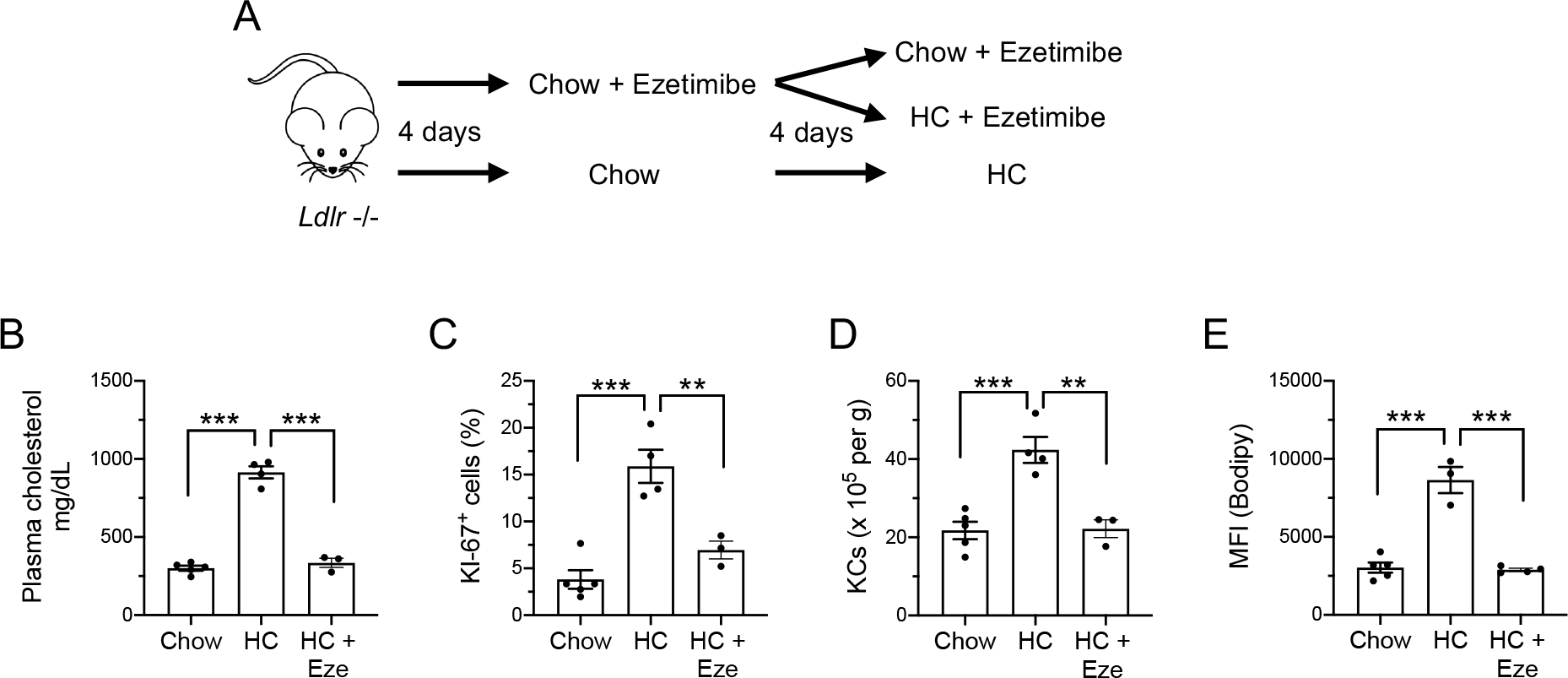
Hypercholesterolemia is the only trigger for KCs proliferation, pool expansion and foamy phenotype. (A) Experimental plan for ezetimibe treatment to block intestinal cholesterol absorption. (B) Feeding ezetimibe to *Ldlr* −/− male mice fed HC diet blocked plasma cholesterol elevation, resulting in (C) no increased proliferation (as determined by KI-67 staining) (D), no increased pool density, and (E) no lipid loading (bodipy) of KCs. (n=4, 4 and 3 for chow-, HC- and HC+eze-fed mice, respectively)

Altogether, we show that EmKCs are actively mobilized during hypercholesterolemia and accumulate large amounts of cellular cholesterol, notably through the scavenging of modified forms of cholesterol-rich LDLs from the bloodstream.

### Monocyte-derived KC with reduced cholesterol-loading sustain the KC pool that contracts during prolonged exposure to hypercholesterolemia

We next evaluated the consequences of prolonged exposure to hypercholesterolemia on KCs. Female *Ldlr*^-/-^ mice were fed the HC diet for up to 3 weeks and analyzed at different time points during this period. After the initial expansion of the KC pool observed at day 4, KC numbers then dropped and stabilized at an intermediate level (Fig. 4A). As mentioned above, previous works revealed that TIMD4 expression is essentially absent on monocyte-derived KCs^10,21^. Here, we observed that the percentage of CLEC2^+^ TIMD4^−^ cells among KCs significantly raised after 8 days of HC diet feeding (Fig. 4B). By day 21, approximately 50% of KCs were TIMD4^−^ (Fig. 4C), suggesting that MoKCs were generated to sustain the KC pool. Absolute cell quantification during this time course analysis clearly showed the progressive engraftment of MoKCs (Fig. 4D). We next generated *Ccr2*^-/-^ x *Ldlr*^-/-^ mice, in which circulating Ly-6C^hi^ monocyte numbers are markedly reduced^30^, and showed that CLEC2^+^ TIMD4^−^ MoKCs were not observed in these animals when subjected to hypercholesterolemia for 3 weeks (Extended data Fig.3A). This confirmed the monocytic origin of MoKCs. After the short expansion phase already described above, we found that EmKCs decreased overtime, returning to steady state levels after 3 weeks of HC diet (Fig. 4D). We thus quantified hepatic *Csf1* mRNA expression levels to determine whether the decrease in EmKCs could result from diminished maintenance signals. *Csf1* mRNA expression progressively increased till day 8 and was then slightly reduced, remaining however elevated by 2-fold as compared to the control chow condition (Fig. 4E). Thus, the decrease in EmKCs numbers after day 4 is unlikely to be the consequence of reduced CSF1 amounts available between day 4 and 8 as *Csf1* mRNA expression kept rising during this time frame. As previously noticed in the context of NASH^10^, MoKCs also presented with a 2-3-fold higher proliferative rate than EmKCs after 3 weeks of HC diet (Fig. 4F). Thus, increased proliferation could confer an advantage to MoKCs as compared to their embryonically-derived counterparts in colonizing the KC niche in the hypercholesterolemic environment. We then asked whether MoKCs generation kinetics would differ between females and males. Thus, changes in KCs homeostasis upon prolonged hypercholesterolemia were also evaluated in male *Ldlr*^-/-^ mice (Extended data Fig. 3B). The contraction of the EmKC pool that occurs after the early proliferative phase was less abrupt in males and only started after day 8 (Extended data Fig. 3C). In addition, MoKCs engraftment started only after day 8 and their proportion among the KC pool after 3 weeks of diet was twice as less than that seen in females (Extended data Fig. 3D). Overall, the results obtained were comparable to those obtained in females, albeit slight differences were noticeable. Having characterized changes occurring to the KC population, we wondered whether inflammatory monocyte-derived macrophages (MoDMacs) were also generated in the liver upon hypercholesterolemia. To do so, we assessed whether we could identify CD64^+^ CLEC2^−^ macrophages by flow cytometry as previously described^10^. While Ly-6C^+^ monocytes increased in the liver after 3 weeks of HC diet, very few MoDMacs could be found (Extended data, Fig.3E). Thus, during prolonged exposure to hypercholesterolemia, monocytes are progressively recruited to the liver and differentiate into MoKCs, but only marginally as inflammatory MoDMacs.

**Fig. 4.**
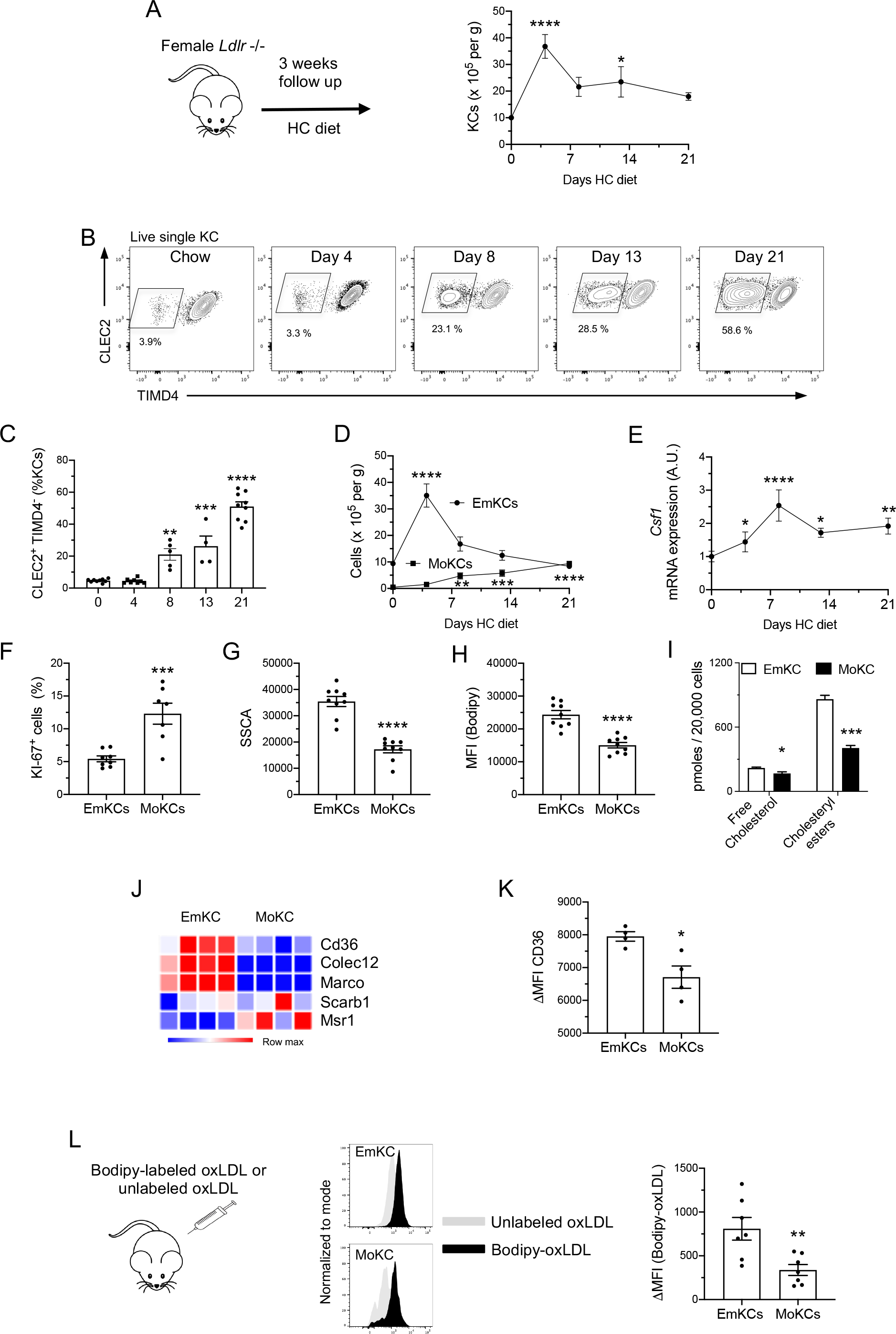
Long term exposure to hypercholesterolemia generates monocyte-derived KCs with reduced lipid-loading capacity. (A) Total KC numbers were determined by flow cytometry before and 4, 8, 13 and 21 days after induction of hypercholesterolemia. (B) Representative flow cytometry analysis showing the progressive increase of CLEC2+TIMD4-KCs with prolonged exposure of the mice to hypercholesterolemia. (C) Frequency of CLEC2 +TIMD4-KCs among KCs (n=9, 8, 5, 4 and 8 mice for experimental points 0, 4, 8,13, and 21 days). (D) EmKCs and MoKCs numbers determined by flow cytometry before (n=9) and at 4 (n=8), 8(n=5), 13(n=4) and 21 (n=8) days after induction of hypercholesterolemia. Indicated p values correspond to significant statistical differences to day 0. (E) Fold-change in *Csf1* gene expression (normalized to day 0) in livers of *Ldlr* −/− mice fed HC diet for the indicated days (n=9, 8, 5, 4 and 8 mice for day 0, 4, 7, 13 and 21, respectively). Indicated p values correspond to significant statistical differences to day 0. (F) Percentage of KI-67^+^ cells among EmKCs and MoKCs in HC-fed female *Ldlr* −/− mice for 3 weeks. (G) Granularity and (H) lipid content (bodipy) of EmKCs and MoKCs of mice fed HC diet for 3 weeks. (I) Lipidomic analysis of sorted EmKCs and MoKCs of mice fed HC diet for 3 weeks. (J) Heatmap depicting the level of mRNA expression of scavenger receptors determined by qPCR on sorted EmKCs and MoKCs from *Ldlr* −/− mice after 3 weeks of HC diet. (K) CD36 mean fluorescence intensity (ΔMFI = CD36 MFI minus MFI of non-stained cells in the corresponding fluorescent channel) determined by flow cytometry on EmKCs and MoKCs at 3 weeks of HC diet (n=4 female mice). (L) Histograms showing MFI in the FITC (bodipy) channel for EmKCs and MoKCs after *in vivo* injection of oxidized-LDL labeled with bodipy or not. The ΔMFI between the conditions is provided for both EmKCs and MoKCs (n=7 mice).

Next, we addressed whether MoKCs properties differed from EmKCs fed the HC diet for 3 weeks. We noticed that MoKCs exhibited reduced granularity (Fig. 4G) and intracellular lipid content (Fig. 4H) as compared to EmKCs in female mice. Similar observations were also made in males (Extended data Fig. 3F and Fig.3G). A reduction in both free cholesterol and all the CE species quantified accounted for the diminished lipid loading observed in MoKCs (Fig. 4I and Extended data Fig. 3H). In addition, RT-qPCR analysis showed decreased expression of the scavenger receptors *Cd36, Colec12* and *Marco*, but unchanged expression for *Scarb1* and *Msr1*, in MoKCs as compared to EmKCs (Fig. 4J). Lower cell surface expression of CD36 protein in MoKCs was confirmed using flow cytometry (Fig. 4K). Thus, we sought to functionally assess whether decreased expression of these scavenger receptors could impact MoKCs ability to uptake modified-lipoproteins. Bodipy-labeled copper-oxidized LDL (oxLDL) were prepared and injected intravenously to evaluate their uptake by KC subsets *in vivo* using flow cytometry. We observed that both EmKCs and MoKCs phagocytized bodipy-labeled oxLDL (Fig. 4L). Nevertheless, bodipy-labeled oxLDL uptake was significantly lower in MoKCs than EmKCs, suggesting that MoKCs have a reduced capacity to scavenge modified-lipoproteins.

Altogether, these results show that hypercholesterolemia triggers a specific adaptive response of KCs with two phases. Following a major EmKCs pool expansion in response to the early induction of hypercholesterolemia, the KC pool is then remodeled with a marked reduction in the number of heavily cholesterol-loaded EmKCs and the engraftment of MoKCs displaying a decreased capacity to scavenge modified-LDLs.

### Protection of EmKCs from apoptosis diminished MoKCs engraftment during hypercholesterolemia

Mitochondrial dysfunction and endoplasmic reticulum stress are important triggers of macrophage foam cell apoptosis^31–34^. In particular, modified forms of LDL and accumulation of cellular free cholesterol can induce the production of mitochondrial reactive oxygen species (ROS) that generate cellular oxidative stress and damage^32,34,35^. We thus sought to determine whether the foamy KCs observed upon hypercholesterolemia underwent mitochondrial stress. KCs were stained with the mitochondria-specific ROS indicator Mitosox and analyzed by flow cytometry. Mitosox fluorescence in EmKCs raised by 1.6- and more than 3-fold when *Ldlr*^-/-^ mice were fed the HC diet for 4 and 21 days, respectively (Fig. 5A), suggesting progressive accumulation of mitochondrial superoxide in EmKCs. At 21 days, mitochondrial ROS detection in MoKCs demonstrated significantly less signal than that measured in EmKCs (Fig. 5B), which might be linked to the lower amount of intracellular free cholesterol measured in MoKCs (Fig.4I).

**Fig. 5.**
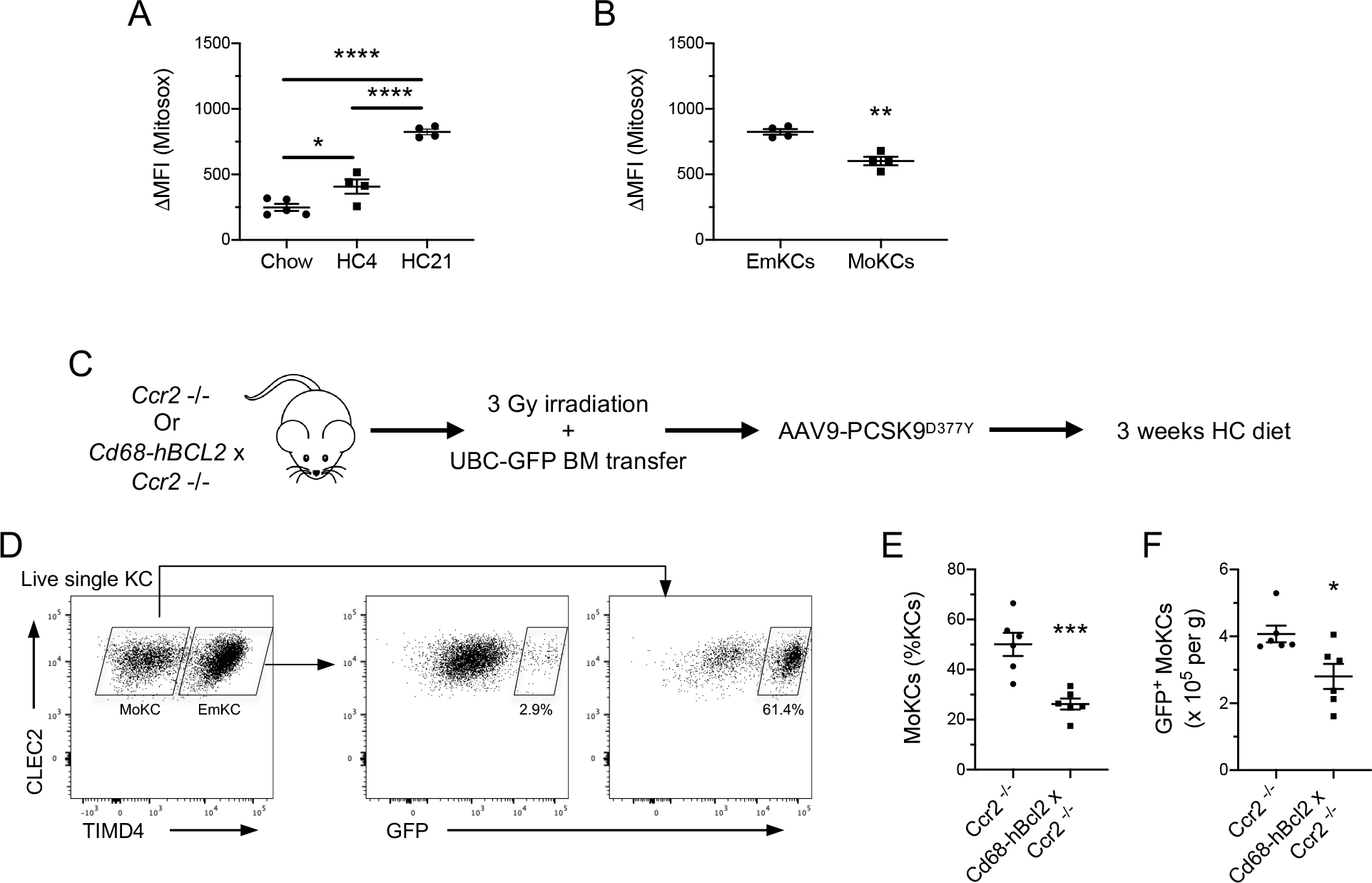
Protection of EmKCs from mitochondrial apoptosis limits MoKCs generation during hypercholesterolemia. (A) Generation of mitochondrial ROS determined by mitosox staining in EmKCs of *Ldlr* −/− female mice fed chow or HC diet for 4 or 21 days (n=4 mice in all conditions). (B) Comparative mitosox staining between EmKCs and MoKCs of *Ldlr* −/− female mice fed HC diet 21 days (n=4 mice). (C) Experimental strategy to track the fate of monocytes in hypercholesterolemic chimeric mice with apoptosis-resistant EmKCs. (D) Representative flow cytometry analysis showing the contribution of monocyte-derived (GFP^+^) cells to EmKCs and MoKCs. (E) Frequency of MoKCs and (F) absolute numbers of GFP^+^ MoKCs in female *Ccr2-/-* (n=6 mice) and Cd68-*hBcl2* x *Ccr2* −/− (n=6 mice) chimeras fed the HC diet for 3 weeks.

We then wondered whether protecting EmKCs from mitochondrial stress would influence MoKCs generation. We thus used *Cd68-hBCL2*^36^ mice exhibiting enforced expression of the mitochondrial anti-apoptotic protein BCL2 in the myeloid lineage, including KCs. *Cd68-hBCL2* mice crossed in a *Ccr2*^-/-^ background were submitted to low-dose irradiation (3 Gy) and subsequently transplanted with bone marrow (BM) cells expressing the green fluorescent protein (GFP). BM^GFP^-transplanted *Ccr2*^-/-^ mice served as controls. In this model, the fate of bone marrow-derived cells can be traced thanks to GFP expression. These chimeric mice were injected with an adeno-associated virus encoding the gain-of-function form of murine PCSK9^D377Y^. PCSK9^D377Y^-mediated degradation of hepatic LDL receptors^37^ then allows for the induction of hypercholesterolemia when mice are subjected to the HC diet (Fig. 5C). After 3 weeks of HC diet, only a small proportion (<3%) of EmKCs expressed GFP (Fig. 5D) in agreement with their embryonic origin, radioresistance and maintenance independent from circulating monocytes. GFP^+^ EmKCs may correspond to the few MoKCs that acquired TIMD4 expression with time, as previously reported^10,21,38^. By contrast, more than 60% of MoKCs were GFP^+^ and thus derived from GFP^+^ donor cells. The remaining GFP^−^ MoKCs most likely corresponded to the few host-derived MoKCs (5-6% of the KC pool at the steady-state, Fig.4C and Extended data Fig. 3D) that proliferated together with the KC pool early after HC diet feeding. The overall frequency of MoKCs was decreased by 2-fold in *Cd68-hBCL2* x *Ccr2*^−/^-recipient mice (Fig. 5E). This was also observed for absolute numbers of MoKCs, whereas EmKCs numbers were increased (Extended data Fig. 4). Finally, quantification of GFP^+^ MoKCs demonstrated diminished amount in *Cd68-hBCL2* x *Ccr2*^-/-^ recipient mice (Fig. 5F). Altogether, these results lend support to the conclusion that protection of EmKCs from cell death, and potentially as a consequence of mitochondrial oxidative stress, limits the recruitment and engraftment of MoKCs under hypercholesterolemia.

### Reducing EmKCs numbers raises cholesterolemia and accelerates atherosclerosis

We finally sought to investigate the potential consequences of decreasing EmKC numbers during hypercholesterolemia. To this aim, we crossed *Ldlr*^-/-^ mice to *Cd207*-*DTR* animals in which diphteria toxin (DT) administration leads to depletion of EmKCs as previously shown^10^. Mice were injected either with DT or boiled heat-inactivated DT (bDT) and then fed the HC diet for 8 days (Fig. 6A). Total KC numbers were similar in both groups after 8 days of HC diet (Fig. 6B). However, DT administration substantially reshaped the KC pool with diminished EmKC counts and a concomitant increase in MoKCs numbers (Fig. 6C). Thus, while few MoKCs were detectable in the bDT group, MoKCs represented about 60% of the KC pool in DT-treated animals (Fig. 6D). As reported above, recruited MokCs were less lipid-loaded than EmKCs in both bDT and DT-treated mice (Fig. 6E). Importantly, both groups exhibited similar counts of Ly-6C^+^ monocytes and neutrophils (Fig. 6F), confirming previous observations that DT administration did not create chronic inflammation^10,21^. In parallel to the increased intracellular lipid content observed in the KCs of DT-treated animals (Fig. 6E), we found that plasma cholesterol concentrations were significantly more elevated in these mice (Fig. 6G). This effect was attributed to EmKCs depletion and not DT off target activity as DT-treated *Ldlr* ^-/-^ mice fed the HC diet for 8 days did not display elevated cholesterolemia (Extended data Fig. 5A). We then assessed whether similar observations could be done when the HC diet was administered for 8 days to *Cd207-DTR* x *Ldlr*^-/-^ mice that received DT and then underwent a 3 weeks recovery period (Extended data Fig. 5B). This set up allows the KC pool to reconstitute before inducing hypercholesterolemia. In this context, EmKC numbers were also found decreased in DT-treated animals and MoKCs counts markedly increased (Extended data Fig. 5C). Total KC numbers was similar in both conditions (Extended data Fig. 5D) but MoKCs represented 40% of the KC pool in DT-treated mice (Extended data Fig. 5E). Again, no overt sign of inflammation was observed upon DT treatment (Extended data Fig. 5F). In this scenario, we observed that plasma cholesterol levels were also more elevated in DT-treated animals (Extended data Fig. 5G). To evaluate the long-term consequences of EmKCs replacement by MoKCs, *Cd207*-*DTR* x *Ldlr*^-/-^ female mice administered DT or bDT were fed the HC diet for 4 weeks (Fig. 6H). Total KC numbers were slightly diminished in DT-treated mice as compared to bDT-treated controls (Fig. 6I). This was primarily due to a marked decrease in EmKC counts as MoKCs numbers were found comparable to the bDT-treated controls (Fig. 6J). MoKCs represented 60% of the KC pool in bDT-treated *Cd207-DTR* x *Ldlr*^-/-^ female mice (Fig. 6K), consistent with our observations in *Ldlr*^*−/−*^ female mice under prolonged hypercholesterolemia (Fig. 4C). This proportion raised to 84% in DT-treated animals (Fig. 6K). Thus, even after several weeks of hypercholesterolemia EmKCs were unable to reconstitute their pool after DT treatment and remain lower in DT-treated mice. Consistent with our findings after 8 days of HC diet feeding, cholesterolemia was also more elevated in DT-treated mice after 4 weeks of diet (1142 mg/dL vs 912 mg/dL in the bDT group, p=0.007) (Fig. 6L). As the liver plays a key role in regulating plasma cholesterol levels, RNA-Seq was performed on livers of these bDT and DT-treated mice. Transcription factor enrichment analysis (CHEA) revealed that many of the down-regulated genes (FC ≥ 1.3) were putative target genes of RXR, LXR and PPARα (based on the use of the ChIP-seq liver mouse dataset GSE35262^39^) (Fig. 6M), all nuclear receptors known for their pivotal roles in the transcriptional control of lipid metabolism. Liver lipidomic analysis confirmed disturbed cholesterol homeostasis (Fig. 6N). Indeed, DT-treated mice exhibited elevated content of cholesterol biosynthesis intermediates (squalene, lanosterol, lathosterol and desmosterol) (Fig. 6N). Liver free cholesterol and cholesteryl-esters were also increased in DT-treated animals as compared to bDT-treated controls (Fig. 6N). Altogether, this shows that transcriptional control of cholesterol metabolism is rewired in the liver of mice exhibiting decreased EmKCs numbers. These results led us to examine whether these changes in cholesterol homeostasis transposed into increased susceptibility to atherosclerosis. Quantification of atherosclerosis in the aortic sinus revealed a 1.6-fold increase in intimal atherosclerotic lesions in DT-treated *Cd207-DTR* x *Ldlr*^-/-^ female mice (Fig. 6O). This experimental protocol was also conducted in male mice and provided similar observations. Notably, DT-treated *Cd207-DTR* x *Ldlr*^-/-^ males exhibited significantly reduced EmKC numbers and elevated hypercholesterolemia (Extended data Fig. 5H-M) that associated with larger atherosclerotic lesions (Fig. 6O). Thus, loss of EmKCs and their subsequent replacement by MoKCs associates with an altered metabolic adaptation of the liver to hypercholesterolemia that favors elevation in hepatic and plasma cholesterol, and accelerated development of atherosclerosis.

**Fig. 6.**
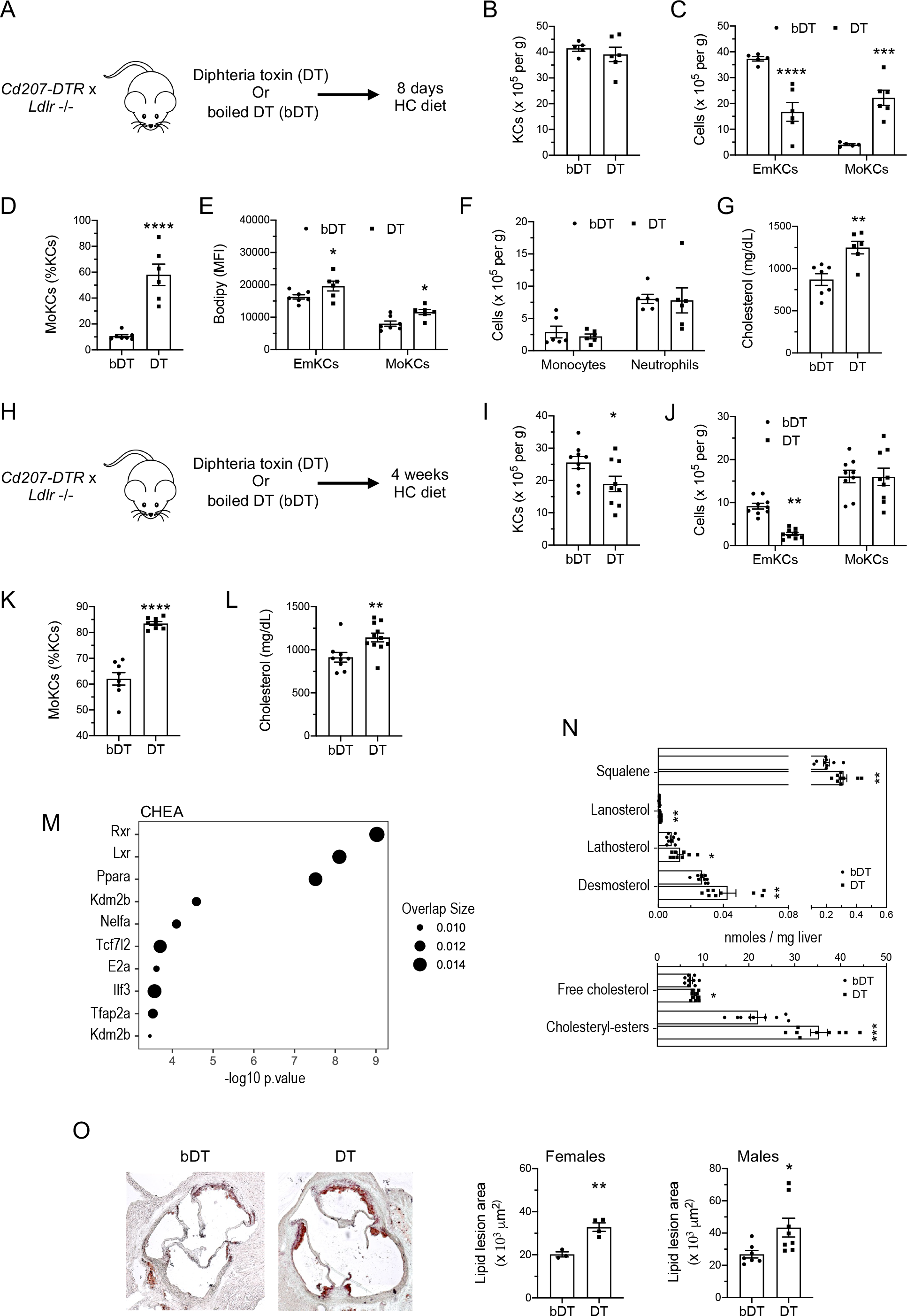
Loss of EmKCs increases hypercholesterolemia and favors atherosclerosis development. (A) Experimental strategy and mouse model used to deplete EmKCs in the context of hypercholesterolemia. (B) Total KC and (C) EmKCs and MoKCs absolute numbers in livers of *Cd207-DTR* x *Ldlr* −/− male mice after 8 days of HC diet and injected either with bDT (n = 7) or DT (n = 6). (D) Percentage of MoKCs, (E) bodipy staining of EmKCs and MoKCs, (F) absolute numbers for liver monocytes and neutrophils, and (G) plasma cholesterol concentrations after 8 days HC diet feeding. (H) Experimental strategy. (I) Total KC and (J) EmKCs and MoKCs absolute numbers in livers of *Cd207-DTR* x *Ldlr* −/− female mice after 4 weeks of HC diet and injected either with bDT (n = 9) or DT (n = 9). (K) Percentage of MoKCs and (L) plasma cholesterol concentrations after 4 weeks HC diet feeding. (M) The list of significantly downregulated genes (RNA-Seq) in livers of DT-treated mice as compared to bDT-treated controls was submitted to the tool Enrichr for transcription factor enrichment analysis (ChEA_2022). (N) Targeted lipidomic analysis performed on livers for cholesterol biosynthesis intermediates, cholesterol and cholesteryl-esters. (O) Representative images of the degree of atherosclerosis in the aortic root area of *Cd207-DTR* x *Ldlr* −/− mice after 4 weeks of HC diet and injected either with bDT or DT. Lipid lesions in the arterial intima were quantified by ORO staining of aortic root sections in female and male mice.

## Discussion

The consequences of hypercholesterolemia on the retention of LDL lipoproteins in focal areas of the arterial tree with their engulfment by macrophages to form lipid-laden foam cells have been studied for decades as an early event in the formation of atherosclerotic lesions.

Here, our work demonstrate that hypercholesterolemia also leads to the generation of heavily cholesterol-loaded Kupffer cells in the liver. We also showed that EmKCs homeostasis is perturbated with a first stage of expansion early after the induction of hypercholesterolemia followed by their impaired maintenance and the engraftment of monocyte-derived KCs while hypercholesterolemia becomes chronic. In addition, the present study lends support for a metabolic function of EmKCs in contributing to cholesterol homeostasis and atherosclerosis development in the context of hypercholesterolemia.

The study of the initial response of EmKCs to elevation of plasma LDL-cholesterol concentration demonstrated a remarkable adaptation of the pool that expanded in both male and female mice. Transcriptomic analysis and KI-67 staining revealed that EmKC expansion was due to increased proliferation. We noticed an increase in liver *Csf1* mRNA expression and a CSF1R inhibitor blunted EmKCs proliferation, strongly supporting a major role for the CSF1-CSF1R axis in this adaptative response of the KC niche. While under steady state conditions hepatic stellate cells (HSCs) have been shown to be the main cellular source of CSF1, liver sinusoidal endothelial cells (LSECs) are also capable to produce CSF1 when KCs are depleted^16^. This suggests that HSCs, and potentially LSECs, were activated in response to hypercholesterolemia to increase hepatic production of CSF1 and sustain EmKCs proliferative response.

After this rapid increase in EmKCs density, we then observed a progressive contraction of the KC pool. The continuous increase in CSF1 expression overtime did not suggest that EmKCs contraction was due to a limitation in the amount of local CSF1 available to sustain EmKCs maintenance. Rather, excessive cholesterol loading of EmKCs may have contributed to decrease their survival capacity. Our lipidomic analyses revealed that, while liver tissue increased its cholesterol content in the form of non-toxic esterified forms of cholesterol, EmKCs accumulated large amount of both free cholesterol and cholesteryl-esters. Excess free cholesterol in macrophages is a potent inducer of their death. Indeed, macrophage free-cholesterol loading has been reported to induce apoptosis, notably by triggering endoplasmic reticulum (ER) stress^40^ but also by increasing mitochondrial oxidative stress and dysfunction^35^. The progressive increase in mitochondrial ROS we observed in EmKCs during HC diet feeding would concur to such scenario. In addition, the involvement of CD36 in EmKCs lipid loading suggested that cholesterol loading is driven by the uptake of modified lipoproteins under hypercholesterolemia. CD36 has been linked to atherosclerosis through its recognition of modified endogenous ligands, including oxidized-LDL^41^. Macrophage uptake of such atherogenic lipoproteins potently induces ER stress and mitochondrial oxidative stress^31–33^, and can even lead to macrophage apoptosis in a CD36-dependent pathway^31^. Thus, under hypercholesterolemic conditions, EmKCs could suffer similar cellular stresses than those operant in lesional macrophages in atherosclerotic plaques.

Loss of EmKCs in depletion models^10,17,21^, in pathological contexts whereby EmKCs are challenged^10–15^ or through genetic invalidation of critical genes for KCs survival^17,42^ have been shown to create niche availability and the engraftment of monocyte-derived KCs. This also occurs in condition of hypercholesterolemia as MoKCs started to progressively emerge when EmKC numbers diminished after the early proliferative phase. The accelerated and stronger loss of EmKCs in females as compared to males also coincided with a more rapid and larger generation of MoKCs in females. It was proposed that niche accessibility and niche availability were the predominant factors for the engraftment of monocytes in a restricted number of niches per organ^43^. The fact that almost no MoKCs were generated in *Ccr2*^-/-^ x *Ldlr*^-/-^ mice strongly suggests that signals of niche accessibility require an operant CCL2-CCR2 axis in this pathological hypercholesterolemic context. Nonetheless, such signaling can be overcome if the niche is brutally emptied as demonstrated in acute depletion models carried on the *Ccr2*^-/-^ background^16^ (an observation that we also verified in hypercholesterolemic DT-treated *Ccr2*^-/-^ x *Cd207-DTR* mice, data not shown). Protecting cholesterol-loaded EmKCs from death through enforced *hBCL2* gene expression, thus limiting niche availability, supported also the proposed niche concept as we observed less recruitment of MoKCs in hypercholesterolemic *Cd68*-*hBCL2* chimeric mice. These data also suggested that loss of cholesterol-loaded EmKCs upon prolonged exposure to hypercholesterolemia was likely linked to mitochondrial-related stress. Such cellular stress could be partially relieved by overexpression of the mitochondrial membrane BCL2 protein that blocks the apoptotic death pathway. With this in mind, reduced cholesterol uptake capacity concomitant to diminished mitochondrial ROS in recruited MoKCs could confer a competitive advantage to these cells over cholesterol-overloaded EmKCs for niche occupancy. Notwithstanding that MoKCs exhibited a higher proliferative state as compared to EmKCs, as previously reported during NASH^10^.

MoKCs have been shown to activate gene expression patterns and display phenotypic characteristics similar to those of EmKCs^16,21^. Nevertheless, specific transcriptomic differences have also been reported between the two cell types^10,13,21^, and functional differences were also highlighted^21,44^. In the context of NASH, we recently revealed that EmKCs were protective over MoKCs to limit liver damage^10^. Here, in the context of hypercholesterolemia, our study adds further support to a beneficial role of EmKCs as compared to MoKCs. Indeed, we observed that both hepatic cholesterol content and circulating concentrations of cholesterol were increased when MoKCs replaced EmKCs in HC-fed and DT-treated *Cd207-DTR* x *Ldlr*^-/-^ mice. On the basis of our transcriptomic analysis, reduced LXR and RXR signaling in livers of DT-treated mice provides a potential explanation for such effects. Indeed, liver-specific deletion of either *Lxra*^45^ or *Rxra*^46^ in mice results in marked cholesteryl-esters accumulation in their livers and high plasma cholesterol levels when challenged with cholesterol-rich diets. LXRs (LXRa and LXRb) form obligate heterodimers with RXR and function as important regulators of cellular sterol homeostasis^47^. LXR / RXR heterodimer can be activated by ligands for either partner, and notably oxysterols and desmosterol for LXRs. Thus, decreased activation of LXR / RXR target genes in the liver of DT-treated mice with less EmKCS and more MoKCs could imply diminished hepatic availability of such ligands (in particular oxysterols as desmosterol is poorly active in hepatocytes^48^). Whether EmKCs contribute to hepatic cholesterol metabolism adjustment to hypercholesterolemia by providing LXR ligands to hepatocytes and whether MoKCs, with a diminished capacity to scavenge circulating oxidized-lipoproteins, might not be as effective as EmKCs to provide such ligands, remain to be established.

Our data also highlight the possibility that the actual development of nanoparticle-based therapeutic strategies to target macrophages within atherosclerotic lesions may also present a great deal of interest to preserve KCs homeostasis and function in the liver of hypercholesterolemic patients. In that respect, it would be highly relevant to assess in preclinical models the beneficial actions on KCs of nanoparticle-mediated delivery of functional miRNA^49,50^ or synthetic LXR agonists^51,52^ that have been shown to promote cholesterol efflux from foamy macrophages, or alternatively, nanoparticle-based approaches to scavenge ROS^53^.

In summary, we have demonstrated that EmKCs are progressively lost and replaced by MoKCs during hypercholesterolemia, which in turn may further exacerbate hypercholesterolemia and atherosclerosis development.

## Methods

### Experimental animals, diets and treatments

All animal procedures were performed in accordance with the guidelines of the Charles Darwin ethics committee on animal experimentation and with the French ministry of agriculture license. This investigation conformed to the European directive 2010/63/EU revising directive 86/609/EEC on the protection of animals used for scientific purposes.

Animals were on a C57BL/6J background. UBC-GFP (C57BL/6-Tg(UBC-GFP)30Scha/J) mice, *Ccr2*−/− (B6.129S4-Ccr2tm1Ifc/J) mice, homozygous *Cd207-DTR* (B6.129S2-Cd207tm3(DTR/GFP)Mal/J) mice and *Ldlr*−/− (B6.129S7-Ldlrtm1Her/J) mice were from the Jackson Laboratory. *Ldlr*−/− were bred in-house to *Ccr2*−/− animals and *Cd207-DTR. Cd68*-*hBCL2* mice previously generated by our team^36^ were bred in-house to *Ccr2*−/− animals. The mice were housed in standard cages at 21 °C and under a 12:12 h light/dark cycle with ad libitum access to water and food. Hypercholesterolemia was induced in *Ldlr*−/− mice by feeding the animals with a chow diet (SAFE A04, Augy, France,) containing 1% cholesterol (HC diet). For studies with ezetimibe, the animals were fed with powdered chow diet (SAFE A04) supplemented or not with 0.005% ezetimibe for 4 days and then switched to powdered chow or HC diets supplemented with 0.005% ezetimibe (Bertin Pharma, France) for 4 days. For CSF1R blocking experiments, 100 mg/kg of PLX3397 or vehicle (0,5% HPMC, 1% Tween 80, 2,5% DMSO) was given to mice by gavage every day from day 0 to day 2 of HC diet feeding. Mice were euthanized and analyzed on day 3.

### Blood, cell-sorted and tissue lipid analyses

Blood samples were collected with heparin-coated capillaries by retro-orbital bleeding in EDTA-containing tubes under isoflurane anesthesia (2% isoflurane/0.2 L O2/min). Plasma samples were stored frozen at −80°C. Plasma total and free cholesterol levels were determined using commercial kits (Diasys). Quantification of cholesterol and cholesteryl ester species in sorted cells or liver tissue was performed by LC-ESI/MS/MS using a Prominence UFLC and a QTrap 4000 mass spectrometer (ICANalytics core facility of the institute of Cardiometabolism and Nutrition (IHU-ICAN, ANR-10-IAHU-05). Briefly, cell pellets or liver tissues resuspended in methanol 70% and homogenized were supplemented with internal standards: CE(18:1)_d7 and cholesterol_d7. Lipids were extracted according to a modified Bligh and Dyer method in methanol/CHCl3 (2:1) and HCl 0.01N. Phase separation was triggered by addition of CHCl3 and H2O. The lower phase was dried and resuspended in LC/MS compatible solvent. Samples were injected to a Ascentis C18 column. Mobile phase A consisted of ACN/H2O (60:40), 10mM ammonium formate, 0.1% formic acid and mobile phase B of ISP/ACN (90:10), 10mM ammonium formate, 0.1% formic acid. Lipid species were detected using scheduled multiple reaction monitoring (sMRM).

### Liver processing and cell suspensions preparation

Mice were euthanized by cervical dislocation. Immediately after, livers were perfused through the portal vein with 1 mL of PBS, followed by 2 mL (1.5ml/min) perfusion with HBSS containing collagenase D (0.2 mU/mL, Sigma). Livers were removed and incubated for 30 min at 37°C under gentle agitation with HBSS containing collagenase D (0.2 mU/mL, Sigma). Cell suspensions were passed through a 100 μm cell strainer before staining. All subsequent procedures were performed on ice.

### Flow cytometry

Antibodies were purchased from BioLegend, Thermo Fisher Scientific, R&D Systems and BD Biosciences. Cell suspensions were pre-incubated 20 mins with anti-mouse CD16/32 antibody (93, biolegend) to block Fc receptors. The following markers and clones were used: CD11c (N418), CD11b (M1/70), CD45 (30-F11), Ly-6C (HK1.4), CD64 (X54-5/7.1), VSIG4 (NLA14), CLEC4F (AF2784), TIMD4 (RMT4-54), CLEC2 (17D9), KI-67 (B56), TCRb (H57-597), CD8a (53-6.7), B220 (RA3-6B2), MHC-II (M5/114.15.2), Nk1.1 (PK136), CD4 (RM4-5), CD36 (BB515), and Ly6G (1A8). Cells were stained with appropriate antibodies for 30 min on ice. Draq7 (BioLegend) was used to exclude dead cells. Intracellular KI-67 staining was performed using the Foxp3 staining kit from Thermo Fisher Scientific. Bodipy 493/503 staining was performed on cells fixed using the Cytofix/CytopermTM kit from BD Biosciences. Mitosox staining was performed before extracellular staining according to manufacturer’s instructions (MitoSOX™ Red Mitochondrial Superoxide Indicator, Thermo Fisher Scientific). To calculate absolute cell counts, a fixed number of nonfluorescent beads (10,000 10-mmpolybead carboxylate microspheres from Polysciences) was added to each tube. The formula number of cells = (number of acquired cells × 10,000) / (number of acquired beads) was used. Cell counts were finally expressed as a number of cells per milligram of tissue. Data were acquired on a BD LSRFortessa flow cytometer (BD Biosciences) and analyzed with FlowJo software (Tree Star). The dimensionality reduction algorithm tSNE (t-Distributed Stochastic Neighbor Embedding) was run using the plugin integrated in FlowJo. KCs cell sorting (EmKCs CD45^+^CLEC2^+^TIMD4^+^CD31- and MoKCs CD45^+^CLEC2^+^TIMD4^−^CD31-) was performed on a BD FACSAria II cell sorter.

### Microscopy

After liver removal, 3-5 mm slices of tissue were fixed by immersion in 4% paraformaldehyde (PFA) for 24h at 4°C, washed in PBS, and further incubated 24h in 30% sucrose. Samples were embedded in Tissue-Tek OCT compound (Sakura Finetek) and frozen using isopentane and liquid nitrogen. 8 μm slices were cut on a cryostat (Leica CM 1900), rehydrated in PBS for 5 mins, and incubated with 0,5% triton and 3% bovine serum albumin for 30 mins at room temperature. Tissue sections were labeled overnight at 4°C in a humid chamber with goat anti-mouse CLEC4F (AF2784, R&D systems) and rat anti-mouse/human CD324 (DECMA-1, BioLegend) antibodies, washed with PBS, and further incubated for 1h at room temperature with cy3 AffiniPure F(ab’)_2_ fragment rabbit anti-goat IgG and alexa fluor 647 AffiniPure donkey anti-rat IgG secondary antibodies (Jackson ImmunoResearch Europe Ltd). Slides were mounted with Vectashield mounting medium with DAPI (Vector Laboratories), imaged with a Zeiss AxioImager M2 microscope (Carl Zeiss) using Zen software. For lipid imaging of liver tissue, *Ldlr*−/− mice fed a chow diet or HC diet for 4 days were anesthetized using Isoflurane and injected intravenously with 5 μg anti-mouse TIM4-APC antibody (RMT4-54, BioLegend). They were then euthanized 5 minutes after injection by cervical dislocation and perfused through the portal vein with PBS and then with 4% ice-cold PFA. Livers were harvested, lobes separated and incubated overnight at 4°C in a fixation/permeabilization buffer (BD Biosciences) diluted 1:4. Tissue samples were washed thoroughly with PBS before imaging. The two-photon laser-scanning microscopy (TPL SM) set-up used was a 7MP (Carl Zeiss) coupled to a Ti: Sapphire Crystal multiphoton laser (ChameleonU, Coherent), which provides 140-fs pulses of near-infrared light, selectively tunable between 680 and 1050 nm and an optical parametric oscillator (OPO-MPX, Coherent) selectively tunable between 1,050 and 1,600 nm. The NLO and the OPO beams were spatially aligned and temporally synchronized using a delay line (Coherent) allowing Coherent anti-stoke Raman Scattering (CARS) imaging approach. The excitation wavelength was 820 nm for the NLO beam and 1070 nm for the OPO beam to detect the vibrational signature of lipid rich structures at a frequency of 2850 cm-1 with an emission wavelength at 665 nm. The system included a set of external nondescanned detectors in reflection with a combination of a LP-600-nm dichroic mirror (DM) followed by a LP-645-nm DM with 624-/40-nm emission filter (EF) and a LP-462-nm DM with 417-/60-nm emission filter (EF), LP-500-nm DM with 480-/40-nm EF, LP-550nm DM with 525-/50-nm and 575/50 nm EFs. Images were performed directly on the whole liver lobe with a water immersion objective, plan apochromat ×20 (numerical aperture = 1). Mask rendering and treatment were done using Imaris software (Bitplane).

### Total body irradiation and bone marrow transplantation

Bone marrow cells were harvested from UBC-GFP donor female mice by gently flushing their femurs. 10 million cells were injected intravenously into 3 Gy irradiated *Cd68-hBCL2* x *Ccr2*−/− or *Ccr2*−/− female mice. A three weeks-recovery period was observed to ensure donor bone marrow engraftment and blood monocytes reconstitution. To ensure induction of hypercholesterolemia, mice were then administered intravenously 10^11^ vector genome copies of a recombinant adeno-associated virus pAAV9-TBG_D377YmPcsk9 encoding the gain-of-function form of murine PCSK9^37^ under the liver specific control of the TBG promoter (virus production by VectorBuilder Inc, USA). Mice were fed 7 days later with HC diet for three weeks before sacrifice. Cell suspensions were prepared from livers and analyzed by flow cytometry as described above. Absolute counts for GFP-expressing cells were normalized to liver monocyte chimerism (94% and ∼84% in *Ccr2*−/− and *Cd68-hBCL2* x *Ccr2*−/− recipients, respectively).

### Diphteria toxin (DT) mediated depletion of KCs in Cd207-DTR mice

Kupffer cells were depleted following two intraperitoneal DT (Sigma) injection (2 × 1μg, 8 to 10 hours apart) in homozygous *Cd207-DTR* X *Ldlr*−/− mice. Heat-inactivated DT (bDT, boiled 25 mins) was used as control.

### In vivo lipid uptake blocking by CD36 neutralizing antibodies

Chow fed *Ldlr*−/− mice were fasted during the day and administered i.v. 50 μg of anti-CD36 monoclonal antibody (MF3, thermoFisher Scientific) or 50 μg of rat IgG2a isotype (thermoFisher Scientific) before given access to HC diet. Antibody injections were repeated the day after, 1 hour before sacrifice.

### In vivo Oxidized-LDL uptake

LDL were isolated from human plasma by sequential ultracentrifugation in the density range of 1.019 < d < 1.063 g/ml, dialyzed against PBS and filter-sterilized. Oxidation was initiated by dialysis of the LDL preparation against a 5μM copper sulfate solution at 37°C. After an overnight incubation, the oxidation reaction was stopped by extensive dialysis against PBS-EDTA 0.1 mM. Changes in the electrophoretic mobility of LDL lipoproteins after oxidation was monitored on agarose gel (hydrogel LIPO + Lp(a) K20, Sebia). Oxidized-LDL (oxLDL) were then incubated with BODIPY 493/503 (4,4-difluoro-1,3,5,7,8-pentaméthyl-4-bora-3a,4a-diaza-s-indacène, ThermoFischer Scientific) (100 μM final) at 37°C for 30 mins and then overnight at 4°C. Free bodipy was removed by passing the oxLDL preparation through a PD-10 column (Pharmacia Biotech, Uppsala, Sweden) using PBS as buffer. Finally, bodipy-labeled oxLDL were concentrated using Spin-X UF concentrator (Corning) to approximately 2 mg protein/mL. 150 μL of the preparation was injected i.v. into *Ldlr*−/− male mice fed 3 weeks HC diet and switched for one week on chow diet to reduce the cellular fluorescence background generated by KCs lipid loading. oxLDL were injected in control mice. Animals were euthanized 1 hour after injection to measure bodipy content of Kupffer cells using flow cytometry.

### qPCR analysis

For gene expression analysis on liver tissue, total RNA preparation was performed using the NucleoSpin RNA Plus kit (MACHEREY-NAGEL). cDNA was synthesized using random hexamer and SuperScript III (Thermo Fisher Scientific). For gene expression analysis on sorted cells, total RNA was prepared from 20,000 cells using the RNeasy Plus micro kit (QIAGEN). RNA was reverse transcribed using the SuperScript VILO cDNA synthesis kit (Thermo Fisher Scientific). Quantitative PCR analyses using Sybr (LightCycler 480 SYBR Green I Master, Roche) were performed using a LightCycler 480 real-time PCR system and dedicated software (Roche). Initial differences in mRNA quantities were controlled using reference mouse genes *18s, Hprt, Rpl13a*, and *Nono*. Primers sequences are available upon request.

### RNA-Seq

Total RNA preparation was performed from 20,000 sorted KCs using the RNeasy Plus Micro Kit (Qiagen) and Nucleospin RNA plus kit for liver samples. cDNA libraries for sorted KCs were generated using Next Ultra II Directional RNA Library Prep Kit for Illumina (New England Biolabs). RNA-Seq libraries were sequenced on an Illumina NovaSeq 6000 (40 million reads per sample). RNA-Seq analysis was completed using the Eoulsan pipeline. The STAR index was used to map raw reads to the genome and data normalization was performed with DESeq2. A LIMMA analysis was conducted to select differentially expressed genes with a 1.3-fold change cutoff between at least two conditions. Adjusted p value for multiple gene testing were used. Annotated genes with a count mean over 100 in at least one condition and a coefficient of variation of more than 0.5 between at least two conditions were retained. GSEA analysis was performed using the Phantasus web platform. The R packages EnhancedVolcano and EnrichR were used for volcano plotting and pathway analyses.

### Data and code availability

The RNA-Seq data generated in this study were deposited under the accession number E-MTAB-12611 and E-MTAB-12744.

### Analysis of aortic lipid lesions

Atherosclerotic lesions were quantified on serial cross sections of the aortic root as previously described^54^. Briefly, mice were anesthetized with isoflurane (2%), euthanized by cervical dislocation and then perfused with PBS. Hearts were collected, fixed with 4% PFA for 30 minutes followed by overnight incubation in phosphate-buffered 20% sucrose solution at 4°C. Hearts were subsequently embedded in Tissue-Tek OCT compound (Sakura Finetek). 10 μm thick sections were cut through the proximal aorta, spanning the three cusps of the aortic valves. 3 sections surrounding the valves (40 μm apart) were fixed and stained with oil red O (ORO, 0.3% in triethylphosphate) for 30 minutes and then counterstained with Mayer hematoxylin for 1 minute. Images were captured using a Zeiss AxioImager M2 microscope and plaque area measured with the AxioVision Zeiss software.

### Statistical analyses

Statistical significance of differences was performed using GraphPad Prism (GraphPad Software). Two-tailed Student’s t test was used to assess the statistical significance of the difference between means of two groups. Experiments were repeated at least twice. Graphs depicted the mean ± SEM. One-way ANOVA and Tukey’s or Dunnett post hoc analyses were used for multiple comparison tests. Statistical significance is represented as follows: *p < 0.05, **p < 0.01, ***p < 0.001, and ****p < 0.0001.

## Acknowledgment

This work was supported by grants to T.H. from the Agence Nationale de la Recherche (ANR-17-CE14-0044, ANR-21-CE14-0067-01), from the Fondation de France and Sorbonne Université Emergence programme.

