## Supplemental figures for "Loss of embryonically-derived Kupffer cells during hypercholesterolemia accelerates atherosclerosis development"

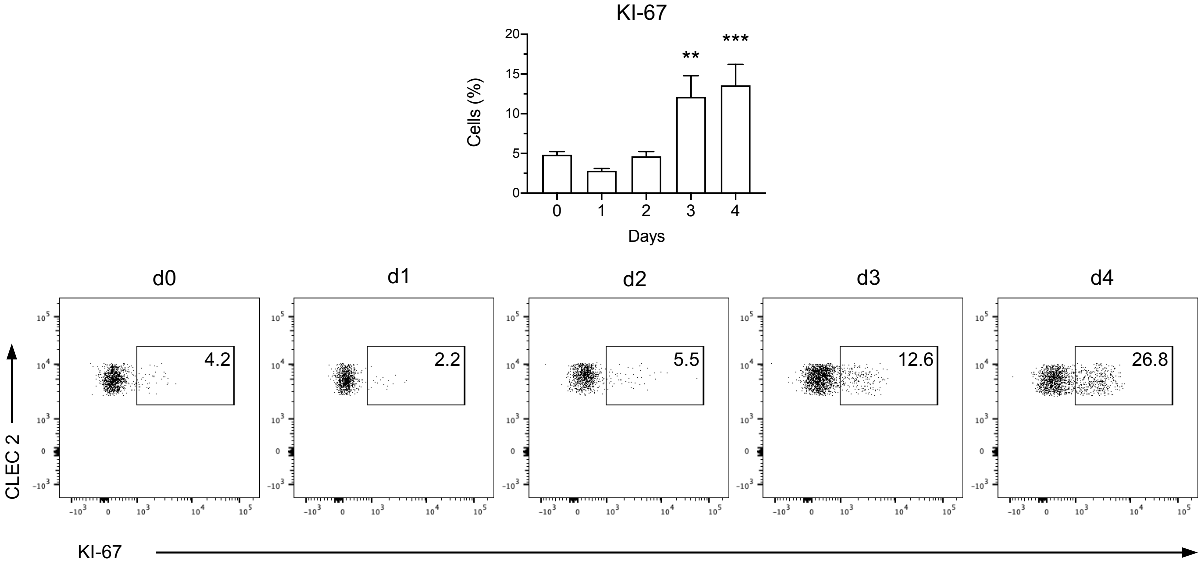


**Extended data Fig. 1**
Percentage of KI-67+ cells among Clec2+ KCs of *Ldlr*-/- mice fed chow or HC diet for 1, 2, 3 or 4 days. Representative FACS plots for each day are shown.

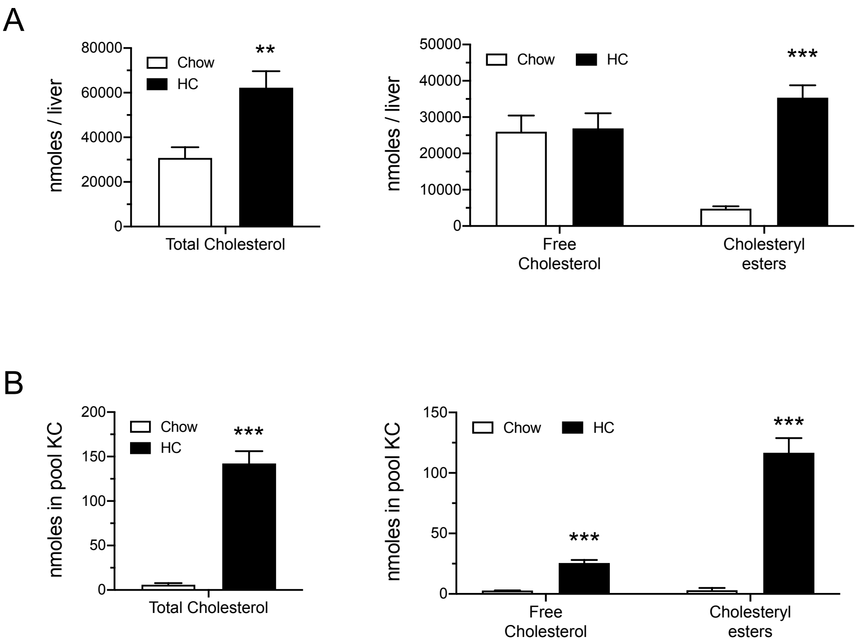


**Extended data Fig. 2**
(A) Calculated amount of total cholesterol and of free and cholesteryl-esters present in the liver of chow- and HC-fed (4 days) *Ldlr-/-* mice. (B) Calculated amount of total cholesterol and of free and cholesteryl-esters present in the KC pool of chow- and HC-fed (4 days) *Ldlr-/-* mice.

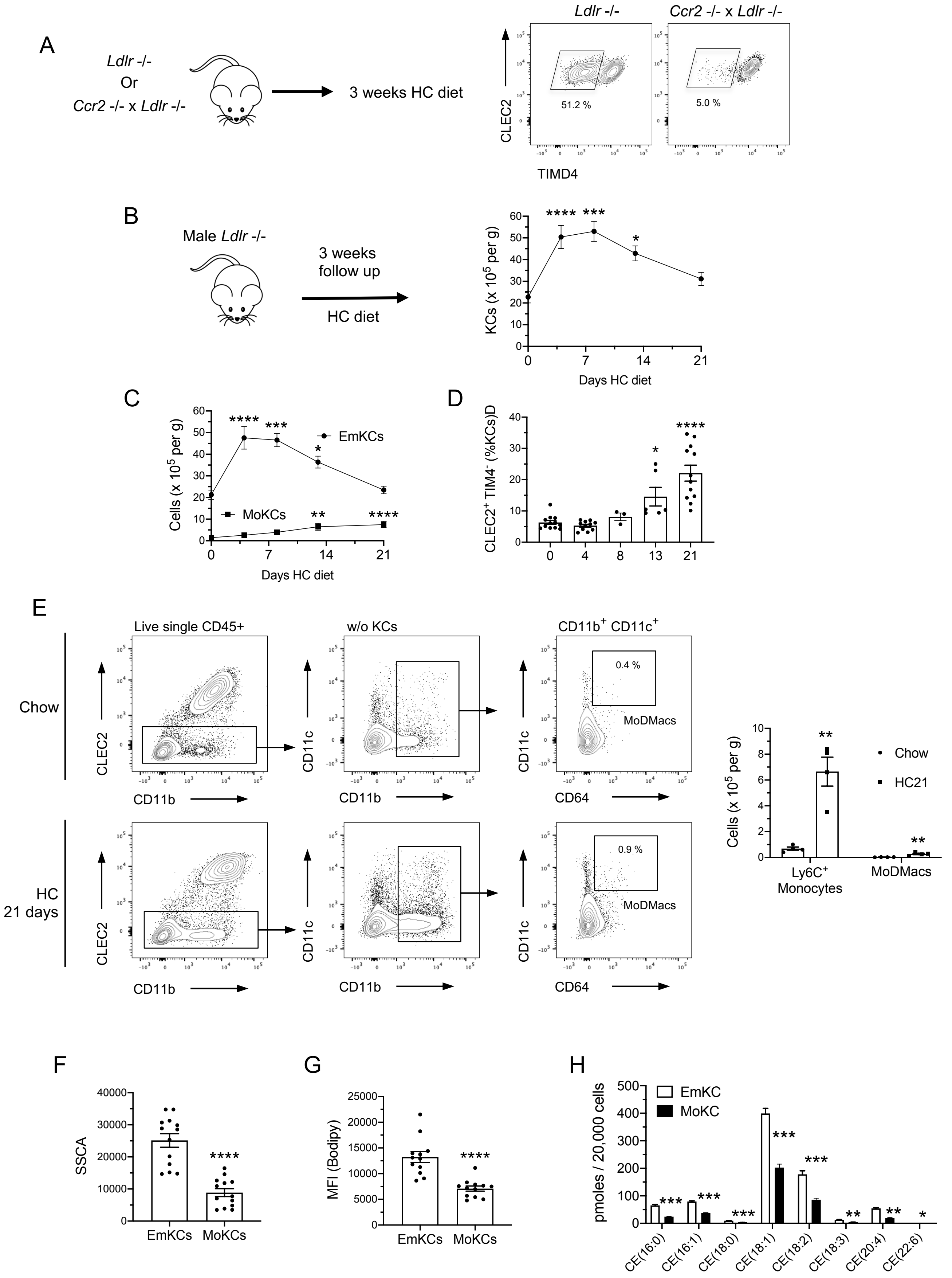


Extended data fig. 3

**Extended data Fig. 3**

(A) Representative FACS plots showing the proportion of TIMD4- cells among CLEC2+KCs in *Ldlr-/-* and *Ccr2-/-* x *Ldlr-/-* female mice subjected to hypercholesterolemia for 3 weeks.

(B) Total KC numbers and (C) EmKCs and MoKCs numbers were determined by flow cytometry before (n=12) and at 4 (n=11), 8(n=4), 13(n=6) and 21 (n=12) days after induction of hypercholesterolemia in male *Ldlr-/-* mice. Indicated p values correspond to significant statistical differences to day 0. (D) Quantification of CLEC2+TIMD4- KCs frequency among KCs.

(E) Representative FACS plots showing the gating strategy used to discriminate MoDMacs in livers of *Ldlr-/-* mice fed chow or HC diet for 21 days. Comparative absolute numbers of Ly6C+ monocytes and MoDMacs found in the livers of the mice under these feeding conditions are shown.

(F) Granularity and (G) lipid content (bodipy) of EmKCs and MoKCs of male mice fed HC diet for 3 weeks. (H) Major cholesteryl-esters species quantified in sorted EmKCs and MoKCs from *Ldlr-/-* mice fed HC diet for 3 weeks.

.

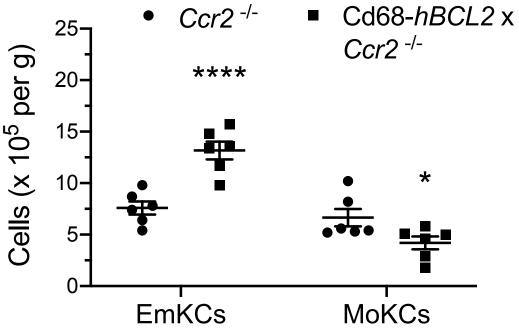


**Extended data Fig. 4**
Absolute EmKC and MoKC numbers measured in *Ccr2-/-* and *Cd68-hBCL2* x *Ccr2-/-* chimeric mice fed HC diet for 3 weeks.

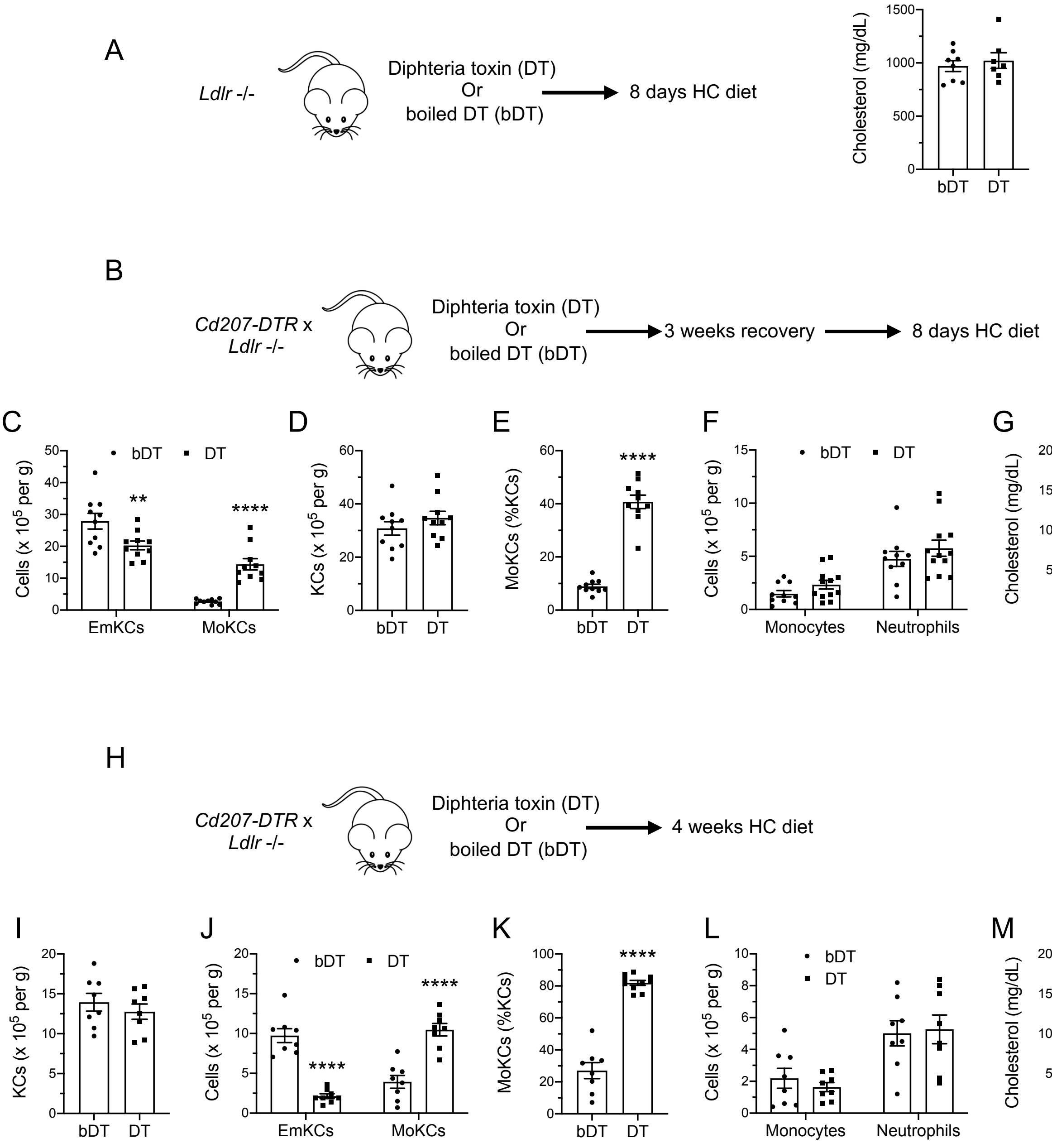


**Extended data Fig. 5**

(A) Absence of plasma cholesterol raising effect in *Ldlr*-/- mice injected with DT and fed HC diet for 8 days as compared to bDT-treated mice.

(B) Experimental strategy with a 3 weeks recovery period post-DT or -bDT injection. (C) EmKCs and MoKCs absolute numbers, and (D) total KC absolute numbers in livers of *Cd207-DTR* x *Ldlr*-/- male mice after 8 days of HC diet and injected either with bDT (n = 10) or DT (n = 12). (E) Percentage of MoKCs, (F) absolute numbers for liver monocytes and neutrophils, and (G) plasma cholesterol concentrations after 8 days of HC diet feeding.

(H) Experimental strategy in *Cd207-DTR* x *Ldlr*-/- male mice. (I) Total KC and (J) EmKCs and MoKCs absolute numbers in livers of *Cd207-DTR* x *Ldlr*-/- male mice after 4 weeks of HC diet and injected either with bDT (n=8) or DT (n=10). (K) Percentage of MoKCs, (L) absolute numbers for liver monocytes and neutrophils, and (M) plasma cholesterol concentrations after 4 weeks HC diet feeding.
